# Lactic Acid Bacteria Dominate Urban Bokashi: A Participatory, Culture-independent Pilot Study of Microbial Diversity and Functional Potential in Household-Scale Food Waste Fermentation

**DOI:** 10.1101/2025.10.16.682835

**Authors:** Katharina Kujala, Veera Kinnunen

## Abstract

In recent years, concerns over declining biodiversity in urban spaces have increased. Urban Bokashi composting (i.e., microaerobic or anaerobic fermentation of food waste indoors) has been suggested as a possibility to promote microbial diversity in the domestic environment. However, studies on microbial communities in household-scale Bokashi and their potential impacts on health and environment are lacking. Thus, the present pilot study investigated microbial communities in different stages of the Bokashi composting process in collaboration with six Bokashi practitioners by looking into physicochemical characteristics as well as microbial community composition (16S amplicon sequencing, 34 samples) and functional potential (shotgun metagenome sequencing, 11 samples). The collective results indicate that i) microbial communities in Bokashi compost differed between stages, but also between households, ii) microbial communities were dominated by lactic acid bacteria like *Lentilactobacillus* or *Lacticaseibacillus*, iii) metabolic pathways for the production of diverse organic acids were detected, iv) application of Bokashi ferment or leachate to soil can supply nutrients and organic acids to promote plant growth but does not substantially affect soil microbial community composition, and v) potentially pathogenic organisms were detected in extremely low abundances. Thus, urban Bokashi is likely not associated with increased health risks and positive impacts are feasible.

## Introduction

In recent years, there has been growing concern over the declining microbial biodiversity of urban spaces and the subsequent disconnect between humans and natural environments (Flandroy *et al*. 2018). In the course of urbanization, the original green spaces have largely been converted into “grey spaces” (United Nations 2004). In many places, natural soil has been replaced by synthetic, impervious materials such as concrete, asphalt or rubber, which has effectively sealed surfaces and impacted thermal and hydrological characteristics of urban spaces. As there is a continuous exchange in microbiota between humans and their environment, low-biodiversity environments can negatively impact the biodiversity of human microbiomes (Hanski *et al*. 2012). Lack of biodiversity in the environment and the human microbiome can in turn have negative effects on the human immune system and lead to a higher prevalence of allergies and autoimmune disorders (Hanski *et al*. 2012; Haahtela 2019). To address this issue, attempts have been made to enhance urban biodiversity by creating green spaces in urban areas such as green walls or roofs and urban gardens. A more recent, promising and increasingly explored option to promote urban microbial biodiversity is the incorporation of soil, rocks, humus and organic matter in public spaces such as playgrounds or parks (Sun *et al*. 2023; Manninen *et al*. 2025). Even enrichment of materials like sand with microbial communities from organic soils has been considered to promote contact of humans with microbial biodiversity (Hui *et al*. 2019).

In addition to larger-scale measures, citizen-driven initiatives can diversify microbial communities and increase exposure to microorganisms in their living spaces such as their homes or their garden. One such practice that has become more popular in recent years in urban settings is Bokashi. Bokashi originates from an East Asian tradition of treating organic waste such as food waste by fermenting it and using the ferment as a soil amendment (Park and DuPonte 2008; Christel 2017). It can be used on the household scale (e.g., (Lew *et al*. 2021)) or in larger settings (Formowitz *et al*. 2007). In contrast to conventional composting, household-scale Bokashi composting is generally practiced indoors in urban settings and can thus lead to a higher exposure of the Bokashi practitioner to microbial communities during the composting process. On the household scale, food waste is collected over the course of several weeks and subsequently fermented microaerobically or anaerobically in closed containers (Kinnunen 2021). To facilitate the fermentation process, the material is typically inoculated with activated Bokashi starter bran which has been prepared with enriched fermenting microorganisms or directly with a solution of fermenting microorganisms. One source of enriched fermenters are the commercially available effective microorganisms (EM®). EM® is a microbial inoculum consisting of about 80 microbial species that was developed in Japan by Teruo Higa (Higa 1991), and is a trademark licensed by EMRO. EM® is used for the preparation of many of the available commercial starter brans. While the detailed composition of EM is kept secret, some of the main species include lactic acid bacteria like *Lactiplantibacillus plantarum, Lacticaseibacillus casei* and *Streptococcus lactis*, photosynthetic purple non-sulfur bacteria like *Rhodopseudomonas palustris* and *Rhodobacter sphaeroides* as well as yeasts like *Saccharomyces cerevisiae* (Xu 2001). Activated Bokashi bran is prepared by mixing enriched fermenting microbial consortia such as EM with wheat or rice bran, molasses and water. This activated bran is subsequently mixed with food waste for fermentation. Alternatively, ready-made Bokashi bran (i.e., prefermented and dried) can be purchased from different suppliers. After a minimum of two weeks of fermentation, the fermented food waste is mixed with soil to complete the process. In addition, leachate is collected from the fermenting Bokashi containers and used for different purposes such as fertilizer (often strongly diluted) or cleaning agent (e.g., to unclog pipes; (Kinnunen 2021)).

Bokashi composting has been surrounded by bold claims for its environmental benefits, such as improving climate smartness and microbial diversity (Fan *et al*. 2018). Although Bokashi practitioners are reporting positive experiences, independent, scientific research demonstrating the environmental impact of the method is still scarce and partly contradictory (see e.g., (Thorslund 2020; Vicente *et al*. 2020; Lew *et al*. 2021; Safwat and Matta 2021). Despite these inconsistent results, there is a shared experiential knowledge among the Bokashi practicing community that using Bokashi as a soil amendment results in a healthy soil by improving soil porosity and water-holding capacity as well as by addition of nutrients, which in turn results in healthier, bigger crops (e.g., (Murillo-Amador *et al*. 2015; Christel 2017; Quiroz and Céspedes 2019)). In addition, bokashi composting has proven to have positive effects especially in odor control and humification (Fan *et al*. 2018). Moreover, vernacular waste fermentation practices, such as bokashi, are also incorporated into urban everyday life in order to remediate toxins and to enrich domestic microbiota and thus enhance wellbeing of ecosystems and inhabitants (Zhang 2021). These holistic health claims are in line with the popularized One Health paradigm which seeks to envision health of people, animals, and environments through the same lens (Evans and Leighton 2014).

While larger-scale nature-based solutions and their impact on health, environment and microbial communities have been studied to some extent, studies on microbial communities in household-scale Bokashi and their potential impacts on health and environment are lacking. Thus, this pilot study set out to characterize microbial communities in Bokashi from six urban practitioners along with collecting information about their experiences with Bokashi composting. Bokashi practitioners were invited to participate in study design and sample collection. The main goals of the study were to i) identify points-of-interest of the Bokashi practitioners and adjust the conducted analyses accordingly, ii) characterize the microbial community composition in different stages of the Bokashi process, iii) investigate the potential for degradation and fermentation processes of actively fermenting Bokashi, iv) identify similarities and differences in Bokashi communities from different households, v) visualize the microbial communities in Bokashi in an easily graspable way for their practitioners, vi) outline potential health benefits and risks associated with household-scale Bokashi, and vii) identify open questions and give recommendation for follow-up studies needed to better understand urban Bokashi composting. To address these goals, Bokashi practitioners provided samples from different stages of their Bokashi process, which were analyzed for microbial community composition and other characteristics. Moreover, participants provided background information on their Bokashi and were able to shape the project focus and data visualization in pre- and post-analysis workshops (Figure 1).

**Figure 1:**
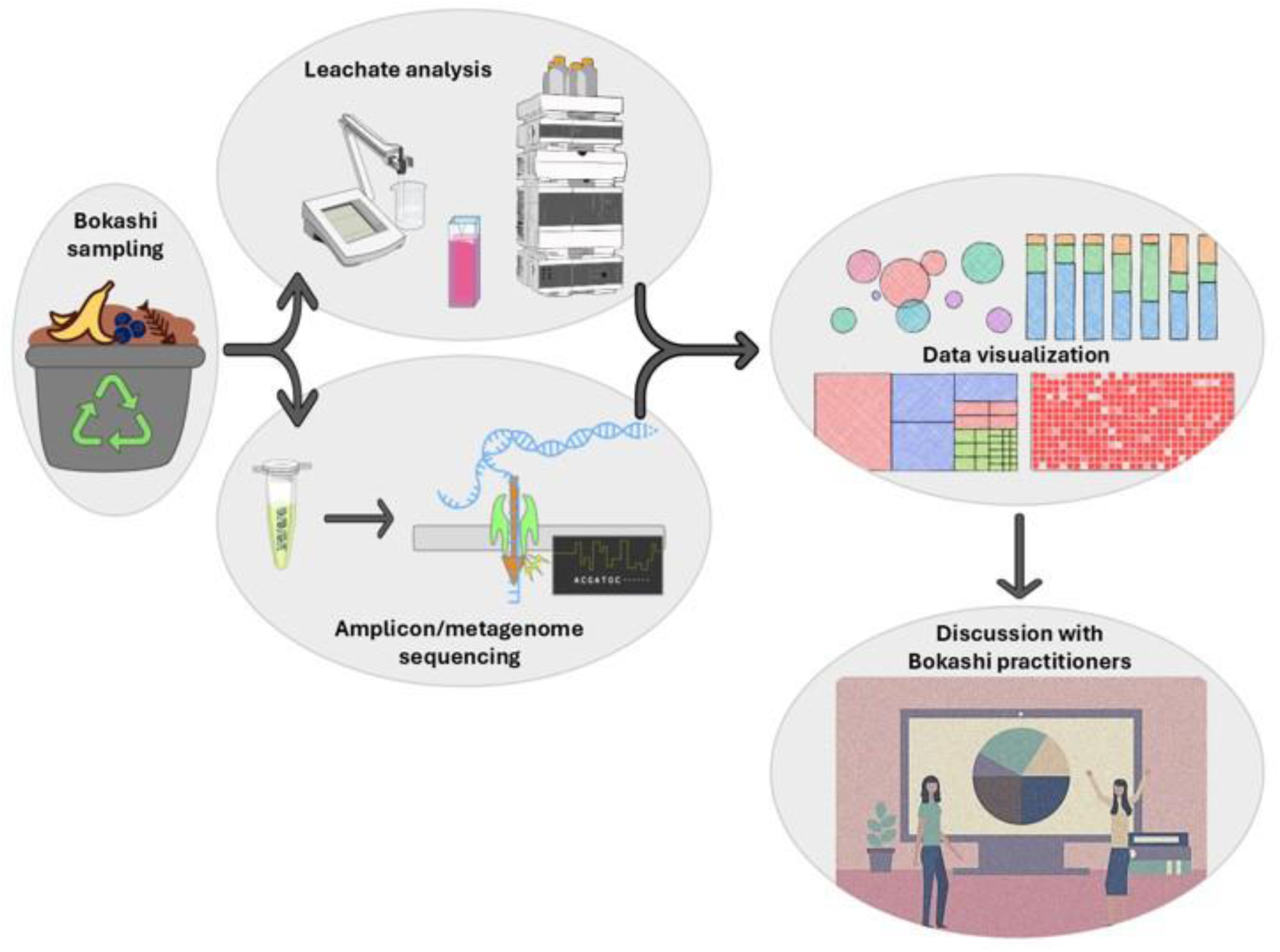
Overview of the project workflow. Samples were collected by six Bokashi practitioners, leachate characteristics and microbial community composition were analyzed and visualized for sharing with the practitioners. The results were discussed with the practitioners to give them insights on the microbial communities in their Bokashi compost and answer questions they brought forward. Icons used to construct this workflow overview were modified from bioicons.com (Agilent_HPLC icon by Sasha Sundstrom, ph-meter and nanopore_sequencing icons by DBCLS, cuvette-pinkdark and microtube-closed-translucent icons by Servier, bubble icon by dbs-sticky, column-stacked-100, heatmap and tree-map icons by dbs-sticky, DNA icon by Kumar) and a graphic designed by Freepick (“People business in office”).

## Material and methods

### Recruitment of Bokashi practitioners

A preliminary study for initial methodology testing was conducted in 2022 with samples provided by one practitioner (more details in “*Citizen sampling of Bokashi*“ below). For the main part of the study, participants were recruited by circulating an open invitation to a face-to-face Bokashi workshop organized at the University of Oulu in May 2023. The invitation was written both in English and in Finnish and it was circulated in two largest social media groups dedicated to Bokashi making (“Bokashi Suomi” and “Bokashi kompostointi (riippumaton)”), on mailing lists of Oulu-based environmental associations as well as mailing lists for University of Oulu researchers. The invitation was addressed to households in the Oulu region which were already practicing Bokashi-composting. The pre-experiment workshop was open for all interested Bokashi practitioners, from which volunteer households were to be selected to participate in sample collection from their Bokashi. Despite a number of people expressing interest in the study, only five Bokashi practitioners attended the face-to-face workshop, so all of them were selected to provide samples for the study. In the beginning of the workshop, the participants were informed about the aims of the study and their rights as research participants, emphasizing that the Bokashi samples will be handled anonymously without any personal identifiers. After the first workshop, two participants withdrew from the study, so two additional Bokashi households were recruited for sample collection through direct contacts.

### Questions brought forward by and interaction with Bokashi practitioners

In the pre-experiment workshop, potential participation in the project was discussed with interested Bokashi practitioners. After giving some initial information about the project, participants were organized into two small groups to discuss what questions intrigued them about Bokashi composting and what issues about Bokashi they would like to have addressed in the experimental phase. The researchers facilitated the discussion and took notes. Practical questions of participants were related to the fermentation process (“What do the microbes produce?”, “What is the pH of the Bokashi/leachate?”, “Is it possible to ferment green waste with EM-liquid without molasses?”, “Do additions like biochar or egg shells affect the process?”), to the use of ferment or leachate (“Does leachate need to be diluted before being used as fertilizer?”, “What type of soil can be used?”, “Is acidity a problem later on?”) and to health implications (“What will stay on my hands if I mix Bokashi/soil without gloves?”, “Are there any proven health benefits?”).

It transpired that Bokashi practitioners wished a rather complete picture of the whole process (from starter to soil) but were especially interested in the analysis of the activated Bokashi starter bran, which is often a commercial product. Based on the participant input and the results of a preliminary study (see next section), four different stages of the Bokashi composting process were selected to be sampled by the Bokashi practitioners (activated Bokashi starter bran, actively fermenting Bokashi, leachate, soil factory).

### Citizen sampling of Bokashi

Two rounds of sampling were conducted. In a preliminary study in 2022, one household (= household 1) sampled actively fermenting Bokashi in different stages (start, after 1 week, after 3 weeks) as well as Bokashi leachate (after 1 week). Based on the outcome of the preliminary study, which showed no major differences between the different stages of actively fermenting Bokashi, and on the input from Bokashi practitioners during the workshop, the protocol was amended to include the following fractions: 1) activated Bokashi starter bran that was used as a Bokashi starter, 2) actively fermenting Bokashi (after 2-5 weeks), 3) Bokashi leachate sampled at the same time, 4) soil collected from the soil factory (i.e., fermented Bokashi mixed with soil; after 2-7 weeks). The duration of each process stage depends on factors such as the type and amount of biowaste produced, the size of the container or space and time constrains and thus varies between different households. As Bokashi practitioners were asked to run their Bokashi process as they typically would, the incubation time for fractions 2-4 provided by the participants was thus equally variable. Samples were taken by the Bokashi practitioners into clean containers (such as glasses or plastic jars of approximately 100-125 ml volume) and frozen in a home freezer. Activated Bokashi starter bran was provided in smaller quantities (∼10g) in plastic bags. Some practitioners provided several samples for stage 2 (5 households) and stage 4 (2 households), either from different fermentation batches or as replicate from the same batch. Samples were then transported by the project team or delivered to the laboratory by the Bokashi practitioners (in styrox boxes with cold elements, transport time < 30 min) and stored at −20°C until further processing. In addition, Bokashi practitioners were asked to provide information about their Bokashi system (e.g., type of food waste that was composted, composting bin type, soil used for soil factory, how long which stage was run).

### Leachate chemical analysis

In the laboratory, Bokashi leachates were filtered (0.2 µm). The leachate pH was analyzed using a handheld pH meter (WTW Sentix). Organic acids were quantified on an Agilent 1200 HPLC system equipped with an Aminex HPX-87H column (300 x 7.8 mm; Biorad) and a UV-visible light diode array detector (detection at 210 nm) using a mobile phase of 4 mM H_3_PO_4_, a column temperature of 60°C and a flow rate of 0.6 mL/min. Organic acid standards with known concentrations of formate, acetate, lactate, propionate and butyrate were used for calibration. Ammonium and phosphate concentrations were determined using a YSI9300 photometer (YSI Incorporated, USA) and the kit/protocols Phot 62 (ammonium) and Phot 28 (phosphate). Nitrate concentrations were determined colorimetrically (Miranda, Espey and Wink 2001).

### DNA extraction, Nanopore 16S amplicon and metagenomic sequencing

DNA was extracted from all samples and a negative extraction control using the ZymoBiomics DNA Miniprep Kit according to the manufacturer’s instructions. Solid materials were used directly (∼250 mg), while leachates were centrifuged for 10 minutes at 14,000 *g* to pellet solid materials (2 mL) and the supernatant was discarded. 16S rRNA genes were amplified using the primer set 515F (5’-GTGYCAGCMGCCGCGGTAA; (Parada, Needham and Fuhrman 2016)) - 926R (5’-CCGYCAATTYMTTTRAGTTT; (Quince *et al*. 2011)). The primers were preceded by adapters (5’-TTTCTGTTGGTGCTGATATTGC-515F, 5’-ACTTGCCTGTCGCTCTATCTTC-926R) to allow for the attachment of barcodes in a second round of PCR using the Oxford Nanopore Technologies (ONT) 96 PCR barcoding extension (EXP-PBC096). The first round of PCR consisted of initial denaturation (5 mins at 95°C), followed by 35 cycles (45 sec at 95°C, 45 sec at 50°C, 90 sec at 68°C) and a final elongation (5 mins at 68°C), while the second round of PCR consisted of initial denaturation (5 mins at 95°C), followed by 15 cycles (45 sec at 95°C, 45 sec at 62°C, 90 sec at 68°C) and a final elongation (5 mins at 68°C). After the second PCR, PCR products were pooled and purified using 1x AmpureXP magnetic beads (Beckman Coulter) according to the manufacturer’s instructions. Purified PCR products were end-prepped and sequencing adapters were ligated according to the ONT protocol. Amplicons were sequenced on a MinION using a Flongle R10.4.1 flow cell as given in the ONT protocol.

Shotgun metagenomes were sequenced only from DNA extracts of actively fermenting Bokashi (11 samples). Due to low sample amount, the rapid PCR barcoding kit (SQK-RPB114.24) was chosen. Input DNA was tagmented which generated DNA fragments of on average 3.5kb (Fig. S1) in size flanked by adapters on both ends. Tagmented DNA was then amplified for 14 cycles using the kit’s barcoded primers and processed as described in the ONT protocol. Metagenomes were sequenced on a MinION using a R10.4.1 flow cell.

### Bioinformatics analysis and representation of the results

Basecalling and demultiplexing of raw POD5 data was conducted with Dorado (GitHub - nanoporetech/dorado: Oxford Nanopore’s Basecaller) using the super accurate basecalling model (dna_r10.4.1_e8.2_400bps_sup@v5.2.0) for both amplicon and metagenome sequencing reads. Demultiplexed amplicon sequence reads were filtered to remove reads with an average quality score < 7 and imported into qiime2 (Bolyen *et al*. 2019). For each barcode, forward and reverse primers were initially processed separately due to the mixed orientation of Nanopore reads. Forward/reverse primer sequences were identified and trimmed at the 5’-end of the reads and the reverse complements of the reverse/forward primer sequences were identified and trimmed at the 3’-end of the reads using the q2-cutadapt plugin (Martin 2011). Reads in which primers were lacking at one or both ends were discarded. The sequence orientation of the reverse reads was adjusted using the q2-rescript plugin (Robeson *et al*. 2021), after which forward and reverse reads and ASV tables were merged. Sequences were classified against the greengenes2 database (McDonald *et al*. 2024) using the consensus blast q2-feature-classifier (Camacho *et al*. 2009). Unassigned reads and reads with taxonomy assigned only at domain-level were removed from the ASV table. The ASV table was collapsed at species-level and rarefied to an even sampling depth of 7000 sequences. Shannon diversity (Shannon 1948), Pielou Evenness (Pielou 1966) as well as Bray-Curtis distance (Bray and Curtis 1957) were calculated based on the rarefied species-level table using the q2-diversity plugin.

Read quantities and qualities of metagenomic reads were visualized as histograms and density plots using Nanoplot (De Coster and Rademakers 2023) and the “ggplot2” (Wickham 2016) and “ggExtra” (Attali and Baker 2025) packages in R version 4.4.2 (R Core Team 2024). The highest density of reads was observed at ∼3.5 kb and an average quality score of 20 in all samp les (Fig. S1), while the reads in the negative control where mainly short (∼200 bp) and of low quality (< 10). Metagenomic reads where filtered using SeqKit2 (Shen, Sipos and Zhao 2024), removing any reads with an average quality score < 10. *In silico* translated reads were aligned against the NCBI non-redundant protein database using DIAMOND (Buchfink, Reuter and Drost 2021) for taxonomic and functional annotation and visualized in MEGAN6 (Huson *et al*. 2007, 2016). Reads were assembled into contigs using myloasm, a tool optimized for the assembly of long-read sequences (Shaw, Marin and Li 2025), which yielded 13,336 contigs with an average length of 11,376 bp (1026 – 2,708,123 bp). Reads were mapped onto the contigs using bowtie2 (Langmead and Salzberg 2012) with mapping rates ranging from 81 to 95% as part of the anvi’o pipeline v8 (Eren *et al*. 2020) and binned using CONCOCT v1.1.0 (Alneberg *et al*. 2014), Maxbin2 v2.2.7 (Wu, Simmons and Singer 2016) and MetaBAT 2 v2.15 (Kang *et al*. 2019). DAS Tool v1.1.7 (Sieber *et al*. 2018) was applied to integrate the results of the three binning methods to produce a set of non-redundant bins, and bins with >50% completeness and <10% redundancy were retained as MAGs. MAGs were imported into Kbase (Arkin *et al*. 2018) for taxonomic classification with GTDB-Tk v2.3.2 (Chaumeil *et al*. 2022; Chivian *et al*. 2023) and functional annotation with DRAM v0.1.2 (Shaffer *et al*. 2020). Phylogenetic trees from GTDB-Tk were modified in iTOL v6 (Letunic and Bork 2024). DRAM output was screened for functions and pathways involved in degradation of organic substrates, including degradation of polymers and monomers as well as fermentations. MAGs were moreover inspected for the presence of potential antibiotic resistance genes using the Resistance Gene Identifier (RGI) web tool of the Comprehensive Antibiotic Resistance Database (CARD) (Alcock *et al*. 2023). Strict and loose hits were filtered to only retain hits with identities >30% to known antimicrobial resistance (AMR) genes and grouped based on the drug classes to which they confer resistance. Notably, some genes confer resistance to multiple drug classes and where thus included multiple times. MAGs were classified as high-quality or medium-quality drafts according to the MIMAG criteria (Bowers *et al*. 2018).

To illustrate microbial community diversity and the dominant genera in each stage of the process to the participating Bokashi practitioners, genus-level relative abundances were visualized as Voronoi plots using the “Weighted Treemap” (Jahn *et al*. 2024) package in R. Only genera with a relative abundance greater than 0.01% were included, and the error tolerance for differences between the target and actual cell sizes was set to 5%. The 15 genera with the highest relative abundance were labelled for each plot. Concentrations of organic acids were visualized using the R package “packcircles” (Bedward, Eppstein and Menzel 2024).

Amplicon and metagenomic sequences were inspected for the presence of potential pathogens. As species-level assignment of short 16S amplicons might not be accurate enough to allow identification of pathogens, genera with potentially pathogenic representatives were also considered. Genera/species identified were *Mycobacterium* (*M. avium, M. lepromatosis*, *M. tuberculosis*), *Clostridium* (*C. botulinum, C. tetani, C. butyricum, C. tertium, C. perfringens*), *Clostridioides* (*C. difficile*), *Bacillus* (*B. licheniformis, B. subtilis, B. anthracis, B. cereus, B. mycoides, B. pumilus*), *Enterococcus* (*E. avium, E. faecium, E. casseliflavus, E. gallinarum, E. faecalis*), *Listeria* (*L. ivanovii, L. monocytogenes*), *Streptococcus* (*S. agalactiae, S. dysgalactiae, S. pneumoniae, S. pyogenes, S. sanguinis, S. suis*), *Mycoplasma/Mycoplasmoides*, *Staphylococcus* (*S. aureus, S. haemolyticus, S. saprophyticus*), *Neisseria, Escherichia* (*E. coli, E. marmotae*), *Shigella* (*S. boydii, S. sonnei*), *Klebsiella* (*K. granulomatis, K. michiganensis, K. oxytoca, K. pneumoniae, K. variicola*), *Salmonella* (*S. enterica*), *Yersinia* (*Y. enterocolitica, Y. pestis*), *Pseudomonas* (*P. putida, P. aeruginosa*)*, Acinetobacter* (*A. baumannii, A. junii, A. soli, A. ursingii*), and *Stenotrophomonas* (*S. maltophilia*).

## Results

### Bokashi setup comparison

Activated Bokashi starter brans from three different companies readily available in Finland were used by the Bokashi practitioners. According to the information provided on the suppliers’ website and/or the product label, all starters were prepared with wheat bran, sugar molasses, water, and EM. Two of the products mentioned additional ingredients such as spelt husk, sunflower oil, and lime (Table 1). No information was provided about the bran activation and drying process (e.g., conditions, duration). Two of the products mentioned some of the microorganisms contained, which included *Lactobacillus plantarum* (= *Lactiplantibacillus plantarum*), *Lactobacillus casei* (= *Lactocaseibacillus casei*), *Rhodopseudomonas palustris* and *Saccharomyces cerevisiae* in concentrations of 10^4^ – 10^5^ cfu g^-1^. While it was not explicitly stated, it seemed like the suppliers listed the species and concentrations that were applied for the preparation of the activated Bokashi starter bran rather than referring to the final product.

**Table 1:** Information on activated Bokashi starter brans provided by the supplier.

| Starter bran | Consistency | Recommended use | Ingredients | EM microorganisms mentioned on label |
| --- | --- | --- | --- | --- |
| 1 | dried grits | add 1 tablespoons to surface for 1L food waste | wheat bran, spelt husk, EM, water, sunflower oil, sugar cane molasses, coralline algae lime | *Lactobacillus plantarum*, *Lactobacillus casei*, *Rhodopseudomonas palustis*, *Saccharomyces cerevisiae* |
| 2 | dried grits | add 2 tablespoons to surface after each addition of food waste | wheat bran, water, sugar molasses, EM-1® | *Lactobacillus plantarum*, *Lactobacillus casei*, *Saccharomyces cerevisiae*, *Rhodopseudomonas palustris*, *Rhodospirillum rubrum* |
| 3 | dried grits | mix 1-2 tablespoon with 1L food waste | wheat bran, spelt husk, EM, sugar cane molasses, EM ceramic powder, seaweed lime, sunflower oil | not specified |

All study participants set up their Bokashi in closed buckets equipped with a stopcock for withdrawal of leachate. In all households, the food waste collected included fruits and vegetables/vegetable peels. In addition, some households reported that they had also added bread (household 2), berries (households 2 and 4), fish (household 3), dairy products (household 3), coffee grounds and tea leaves (household 5), porridge (household 6) and meal leftovers (household 6). However, no guarantee can be given for completeness of information, as the collection of food waste was conducted over a period of several weeks, and study participants did not provide an exhaustive list of the contents of their Bokashi containers. For the soil factory, study participants used garden and sandy soil (from their own garden or flowerpots) as well as leftover soil from previous soil factories.

### Physicochemical characteristics of Bokashi leachate

All leachates were acidic with pH ranging from 3.7 to 4.9 and contained large quantities of organic acids (Table 2). In most households, lactate was the dominant organic acid produced with concentrations ranging from 14 to 19 mM (Table 2). Only leachate from household 5 had considerably lower lactate concentrations (< 5 mM). Acetate and propionate were present in similar concentrations in all leachates (9-14 mM and 1.5-2.5 mM, respectively), while butyrate had only accumulated to >5 mM in four of the leachates (households 1, 3, 5, 6). Formate concentrations were rather low in all leachates (< 0.25 mM). Ammonium concentrations in leachate ranged from <10 to 167 µM, while nitrate concentrations ranged from 29 to 860 µM. Phosphate concentrations ranged from 3 to 70 µM.

**Table 2:** Characteristics of Bokashi leachate.

| Household | pH | EC<br>(ms/cm) | Ammonium<br>(μM) | Nitrate<br>(μM) | Phosphate<br>(μM) | Formate<br>(mM) | Acetate<br>(mM) | Propionate<br>(mM) | Butyrate<br>(mM) | Lactate<br>(mM) |
| --- | --- | --- | --- | --- | --- | --- | --- | --- | --- | --- |
| 1 | 4.661 | 13.5 | <10 | 189.5 | 20.7 | 0.20 | 11.8 | 2.3 | 5.0 | 14.3 |
| 2 | 3.778 | 10.9 | 11.1 | 857.6 | 2.7 | n.d.<sup>a</sup> | 12.7 | 1.5 | 0.37 | 13.0 |
| 3 | 3.852 | 15.2 | 111.1 | 44.2 | 69.7 | 0.05 | 14.6 | 2.3 | 6.3 | 17.2 |
| 4 | 3.706 | 13.73 | 111.1 | 595.3 | 6.7 | n.d. | 13.0 | 1.8 | n.d. | 19.4 |
| 5 | 4.909 | 10.63 | 166.7 | 23.3 | 38.7 | 0.15 | 8.9 | 2.5 | 7.1 | 4.6 |
| 6 | 4.335 | 14.89 | 138.9 | 29.1 | 41.9 | 0.21 | 10.3 | 1.9 | 5.8 | 18.5 |

<sup>a</sup> not detected

### Microbial communities during different stages of the Bokashi composting process

16S rRNA gene amplicon sequencing revealed that microbial communities in actively fermenting Bokashi and its leachate were dominated by *Firmicutes* (50-99% relative abundance; Figure 2 A). Proteobacteria were the second most abundant group, accounting for up to 30% of the microbial communities in this stage. However, *Proteobacteria* were only abundant in Bokashi and leachate from households 2, 5 and 6. The dominant genera in actively fermenting Bokashi and its leachate differed between the households: While *Lentilactobacillus* was rather abundant (20-60% relative abundance) in samples from most households (apart from household 5), the abundance of other genera was more individual (Figure 2 B). *Companilactobacillus* had a relative abundance of 12-27% in household 1 but was of low abundance in all other households (<4%). *Lactobacillus* had a high relative abundance (25-30%) in actively fermenting Bokashi and leachate from household 3, while of low abundance (<2%) in all other samples. *Lactiplantibacillus* and *Levilactobacillus* were prominent in samples from households 2 and 4. Households 2 and 5 had many genera below the threshold abundance of 5% in at least one sample. Microbial community composition in Bokashi starter bran was often similar to the composition in actively fermenting Bokashi of the same household (Fig. S2 A), and *Lentilactobacillus* was the dominant genus in starter brans from supplier 1 and 3. Notably, photosynthetic genera like *Rhodopseudomonas, Rhodobacter* and *Rhodospirillum* (which are advertised as a beneficial part of EM) were not detected in any of the analyzed activated Bokashi starter brans (at a sequencing depth of >7000 sequences), indicating that they were not present in high numbers in the used starters. *Rhodopseudomonas* and *Rhodobacter* were detected in some soil factory samples, indicating that their detection could be achieved with the used protocol. It is, however, possible that the used DNA extraction kit worked less well for the activated Bokashi starter brans than for the Bokashi or soil factory samples.

**Figure 2:**
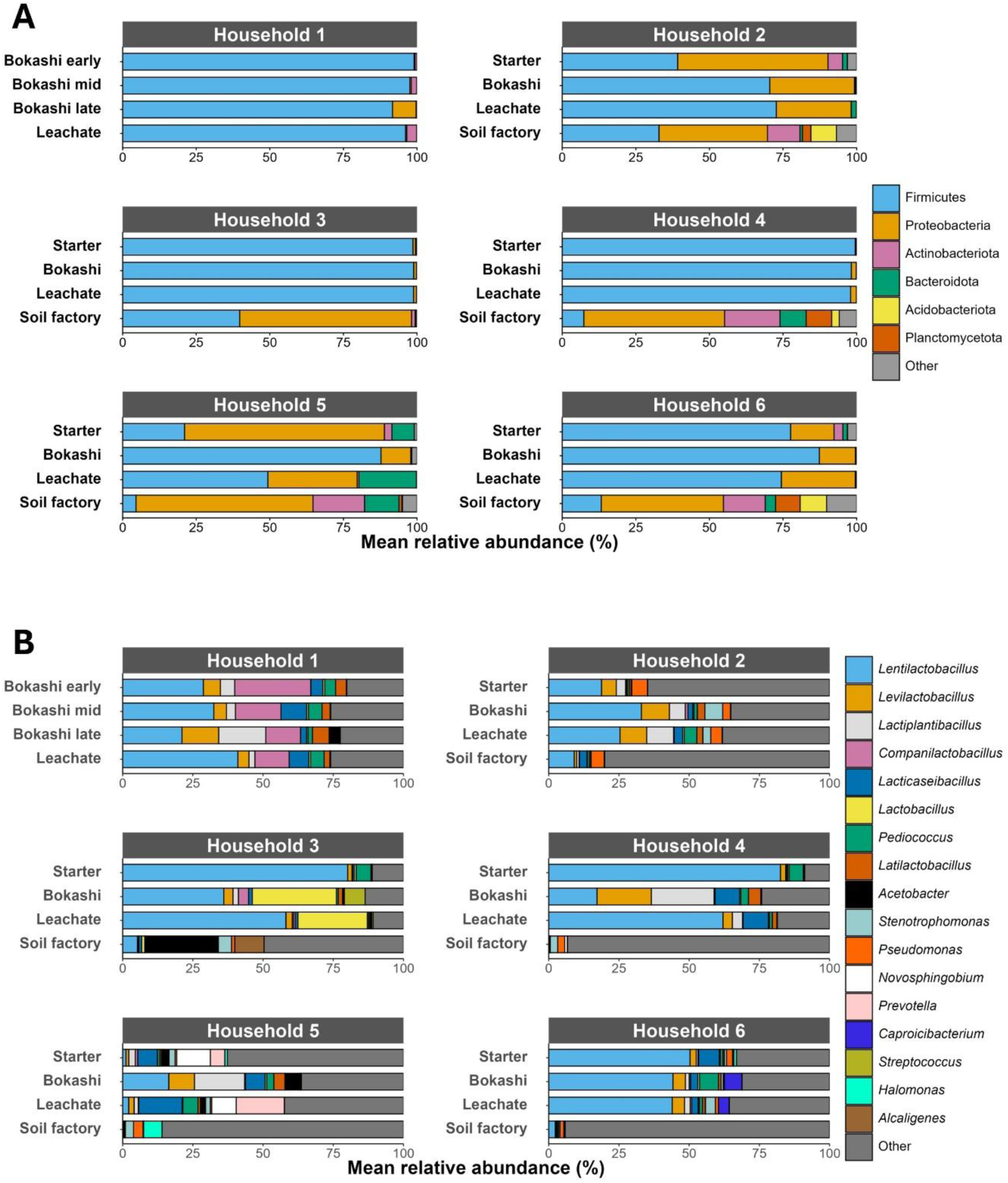
Phylum- (A) and genus-level (B) microbial community composition in different stages of the Bokashi process based on 16S rRNA gene amplicon sequencing. Phyla or genera with a relative abundance < 5% in all individual samples were grouped (“other”). Average relative abundances are presented for cases in which duplicate samples were analyzed (marked with *).

Microbial diversity in soil factories was considerably higher than in the activated Bokashi starter brans, actively fermenting Bokashi or leachate, with a median number of observed species of 1100 (vs. 200-500) and a median Shannon diversity of 8 (vs. 4-5; Figure 3). Microbial communities in soil factories were dominated by Proteobacteria (37-60%; Figure 2 A). In households 2 and 3, Firmicutes were prominent (33-39% relative abundance), while in the other households they were below 10%. Unlike the actively fermenting Bokashi samples, most soil factories were not strongly dominated by one or two genera but had a more even distribution (Figure 2 B), indicating that soil factory microbial communities were likely not determined by the Bokashi ferment they received but rather by the residing soil microbes.

**Figure 3:**
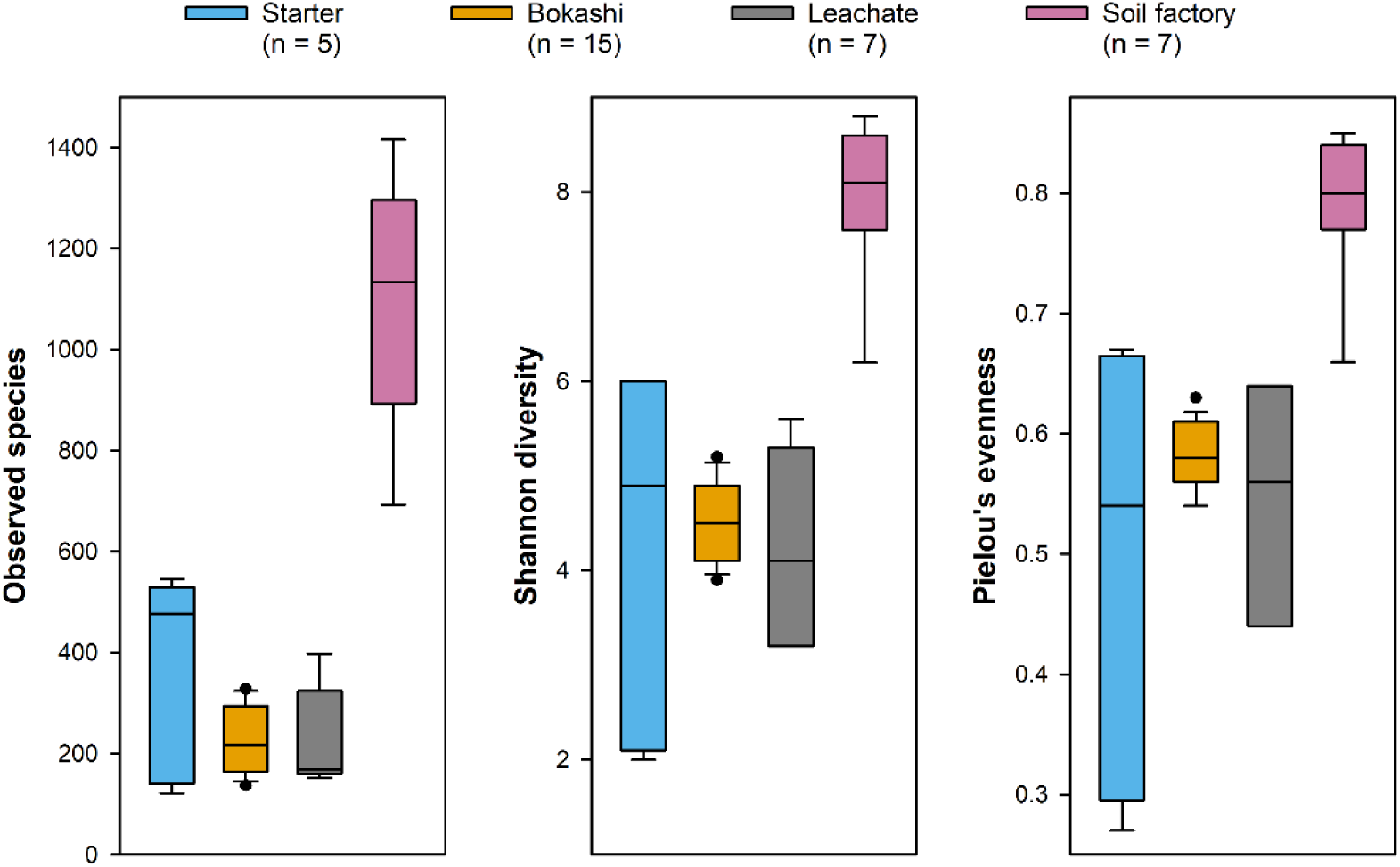
Species-level microbial diversity in different stages of the Bokashi process based on 16S rRNA gene amplicon sequencing. No. of observed species, Shannon diversity and Pielou Evenness diversity indices were calculated based on the rarefied species table using the q2-diversity plugin after rarefication to an even sampling depth of 7000 sequences.

**Figure 4:**
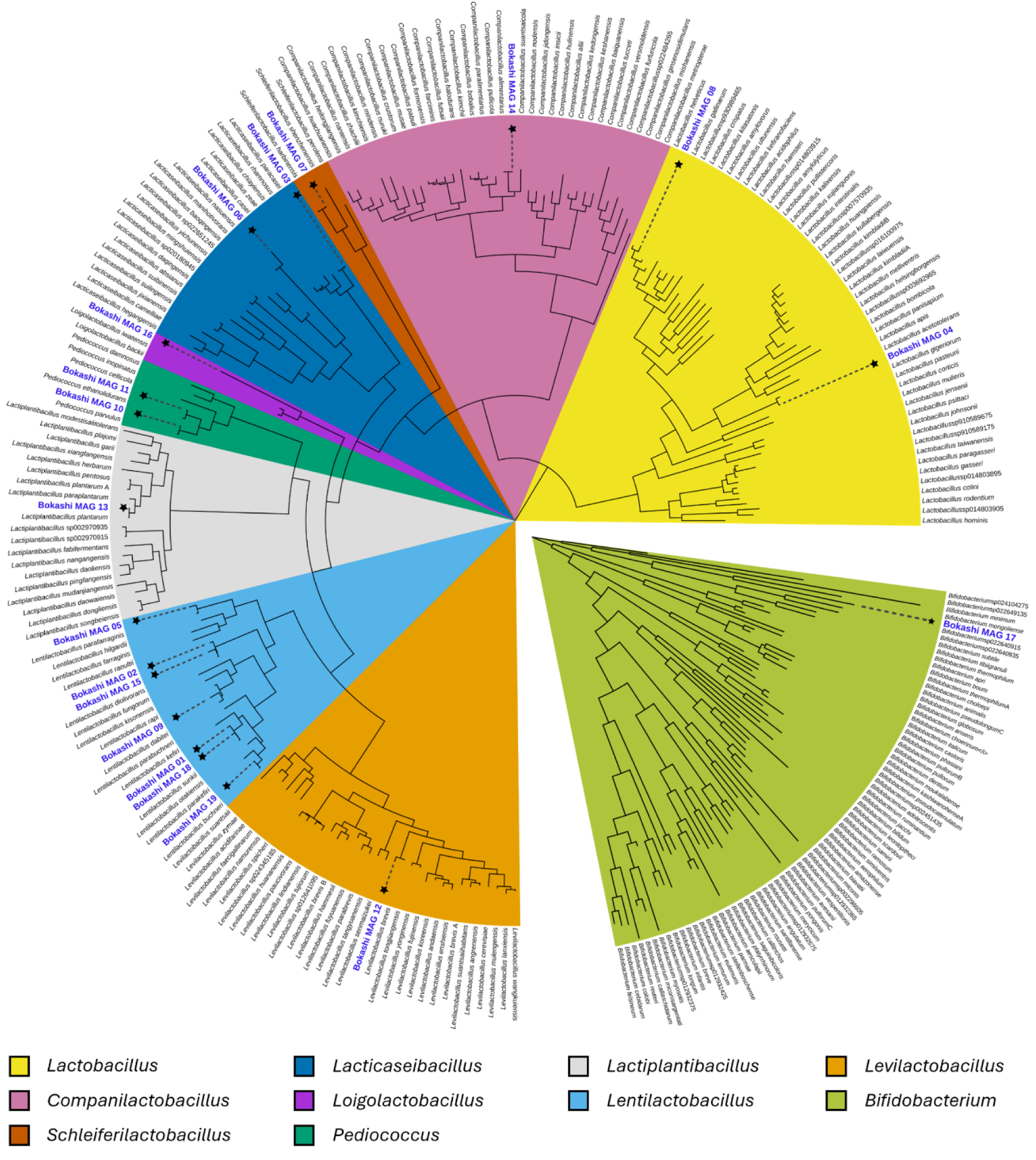
Phylogenomic tree of the 19 metagenome-assembled genomes (MAGs) recovered from actively fermenting Bokashi. MAGs were placed in GTDB species trees for lactic acid bacteria and Bifidobacteria using GTDB-tk Classify in KBase. The tree was imported into iTOL v7 for midpoint rooting and editing. MAGs are indicated by (*) on the branches and bold, blue font.

### Metagenomic analysis of actively fermenting Bokashi

Long-read metagenomes were created from eleven samples of actively fermenting Bokashi. Analysis of unassembled reads mostly replicated the prokaryotic relative abundance patterns and community similarity patters obtained from 16S rRNA gene amplicon sequencing (Fig. S3, Fig. S2 B). A small percentage of metagenomic reads (0.6 to 1%) was affiliated to Eukaryota and viruses (Table S1). Some species like *Saccharomyces cerevisiae* or *Aureobasidium melanogenum* were only detected in one or two samples, while others like *Trichuris trichiura* or *Mortierella* sp. were detected in all samples. The detected viruses were mainly viruses of *Lactobacillus/Lactobacillaceae*. SEED functional annotation revealed the functional potential for organic matter degradation and fermentation processes (Fig. S4).

In each household, actively fermenting Bokashi was typically dominated by one to three MAGs (Fig. 5). The functional potential of MAGs was inspected focusing on pathways and functions important for organic matter degradation as well as for antimicrobial resistance. MAGs encoded pathways for the degradation of glucose as well as enzymes needed for the degradation of complex organic compounds, including beta-glucosidase, beta-galactosidase and arabinosidase. Moreover, fermentative pathways that produce short chain organic acids or alcohols were also encoded, with potential for lactate, acetate and alcohol production being present in nearly all MAGs, while the potential for butyrate, propionate or formate production was detected in only a few MAGs. Hits to potential antibiotic resistance genes were found in most MAGs, with the highest number of hits for genes conveying resistance to fluoroquinolones, macrolides and tetracyclines (Fig. S5). However, the majority of the hits were classified as “loose” by the CARD search tool with only 24 “strict” hits, which mainly were related to resistance to the glycopeptide antibiotic vancomycin (15), disinfecting agents like benzalkonium chloride (5) and fluoroquinolones like norfloxazin and ciprofloxazin. Loose hits, while showing good sequence similarity to known antimicrobial resistance genes, might not actually encode functional proteins involved in antimicrobial resistance.

**Figure 5:**
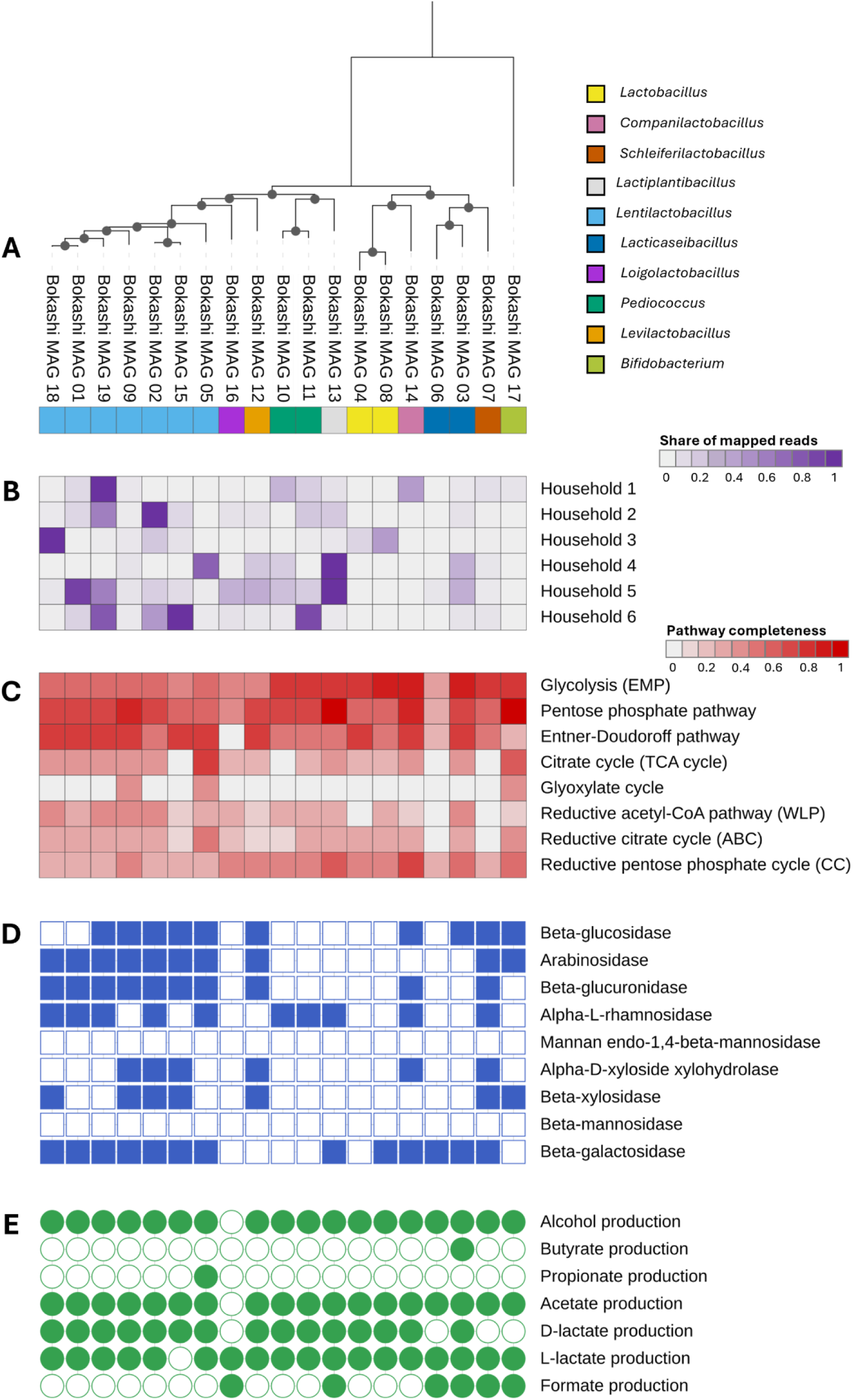
Taxonomic identification (A), relative abundance (B) and metabolic potential (C-E) of metagenome-assembled genomes (MAGs) recovered from actively fermenting Bokashi. In (B) averages of duplicates for households 1-3 and 5-6 are shown. Pathways and functions were identified based on DRAM annotations. Multiple variants of organic acid production pathways provided by DRAM were combined in (E). The tree and heatmaps were generated using iTOL v7.

### Presence of potential pathogens in different stages of Bokashi composting

The relative abundance of potential pathogens was low in all samples. When using classification to the species-level for amplicon sequence data, 0.2 % (0.03-1.1%) were identified as potentially human pathogenic species (Table S3). Most of the species detected were opportunistic and only rarely cause disease (or only in immunocompromised individuals). On average, the three most common potential pathogens were *Klebsiella pneumoniae* (0.034%), *Salmonella enterica* (0.029%), and *Pseudomonas aeruginosa* (0.021%). When using genus-level classification for amplicon sequencing data, 4.7% (0.34-15%) of detected genera in a sample were known to host pathogenic species or strains. On average, the three most common genera with known pathogens were *Stenotrophomonas* (1.2%), *Pseudomonas* (1.0%), and *Klebsiella* (0.50%). However, while these genera host some potentially human-pathogenic species, they also host many non-pathogenic species such as *P. fluorescens, P. stutzeri, K. phyllosphaerae,* or *S. rhizophila.* In the metagenomic reads from actively fermenting Bokashi, detection of potential pathogens was considerably lower: on average only 0.04% and 0.3% of the reads were classified as potentially human-pathogenic species and genera, respectively (Table S3). The most abundant species were *Enterococcus faecalis* (0.026%), *Enterococcus faecium* (0.008%), and *Streptococcus pneumoniae* (0.002%), while the most abundant genera were *Clostridium* (0.29%), *Enterococcus* (0.035%), and *Streptococcus* (0.003%).

### Communicating the results to study participants

For study participants, information cards were generated to illustrate the microbial community composition in different stages and the characteristics of the leachate (Figure 6). These results were shared at a post-experiment workshop, in which the questions brought forward by participants in the previous meeting or after seeing the results were also addressed. Participants were concerned when names of know pathogenic genera appeared in their chart, e.g., *Pseudomonas, Klebsiella* or *Streptococcus*. Therefore, the workshop was used as an opportunity to discuss the diversity of the genera in question, which are metabolically versatile and contain also many non-pathogenic species as well as microbes common in soil and water. Species-level classifications of sequences were not shared with the participants due to potential inaccuracies caused by incomplete coverage of the 16S rRNA genes and potential sequencing errors. The results of the metagenomic analysis were not yet available at the time of the workshop and were thus not discussed. Overall, participants showed great interest in the study results, and some of the questions from the pre-experiment workshop could be answered (e.g., regarding pH or what the Bokashi microbes produce). However, many open questions remained that could not directly be answered with the experimental results and further questions were brought forward during discussions in the post-experiment workshop. E.g., Bokashi practitioners questioned whether the addition of activated starter bran was actually needed, or whether inoculation with leachate or ferment from a previous batch would be enough, and brought forward many practical concerns e.g., related to regulations for composting by local authorities. In hindsight, a second post-experiment workshop might have been needed to better connect with the Bokashi practitioners. Many questions (e.g., related to health impacts of Bokashi) could not be addressed directly due to the study design, but answers might in part be found in the scientific literature or might be addressed in follow-up studies

*Figure 6: Example (household 6) of visualization provided for the Bokashi practitioners in the final workshop. (A) Microbial community composition in each stage, illustrated as Voronoi plots in which the polygon area correlates with the relative abundance of the genera. Top 10 genera are labelled in each plot. (B) Leachate characteristics. Organic acid concentrations are visualized in a bubble plot, in which the bubble size corresponds to the measured concentration in µM. Visualizations for all households are given in the supplementary material. Note that the original version of the visualization used during the workshop were prepared in Finnish language.*

*Figure 7: Example (household 6) of visualization provided for the Bokashi practitioners in the final workshop. (A) Microbial community composition in each stage, illustrated as Voronoi plots in which the polygon area correlates with the relative abundance of the genera. Top 10 genera are labelled in each plot. (B) Leachate characteristics. Organic acid concentrations are visualized in a bubble plot, in which the bubble size corresponds to the measured concentration in µM. Visualizations for all households are given in the supplementary material. Note that the original version of the visualization used during the workshop were prepared in Finnish language.*

**Figure 8:**
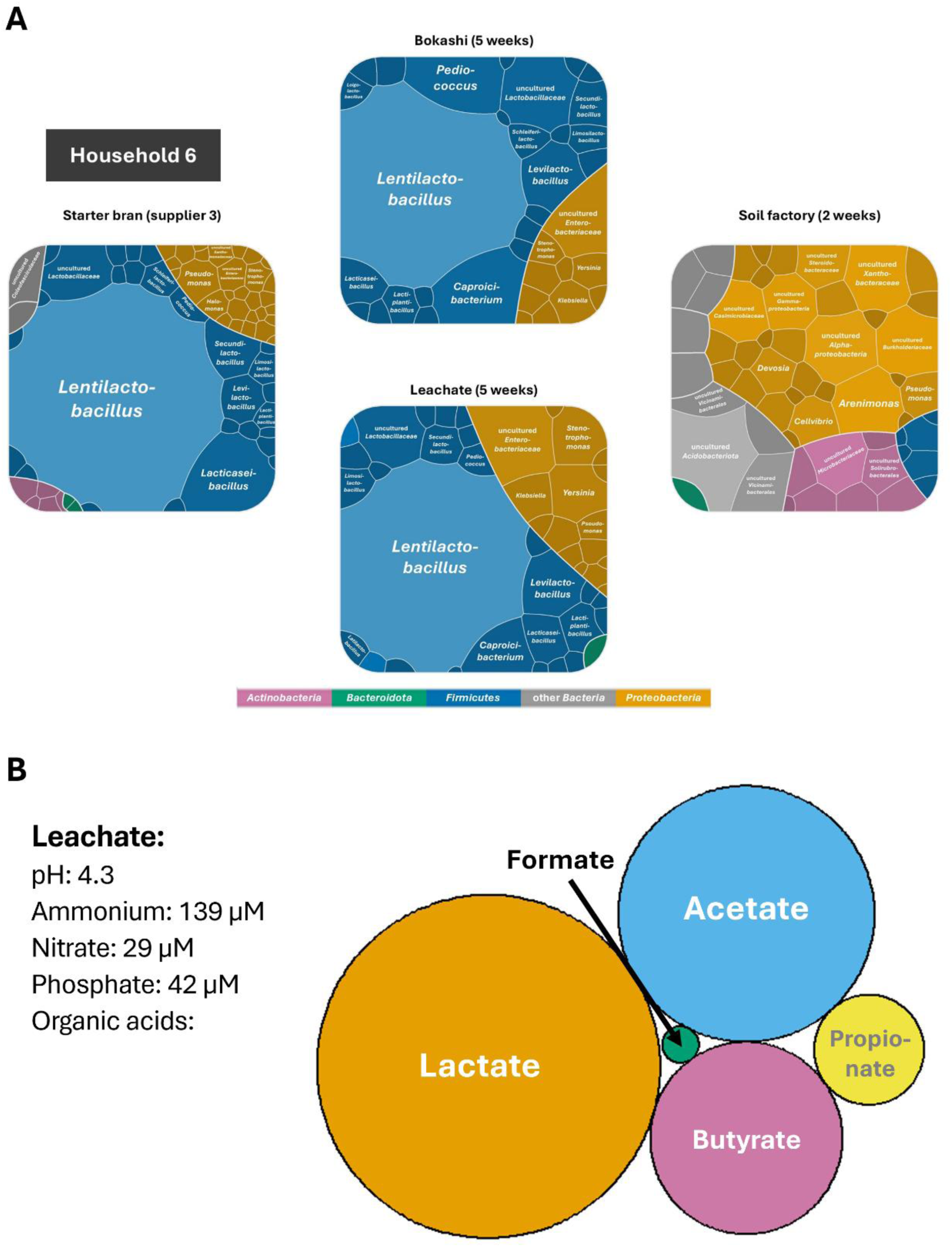
Example (household 6) of visualization provided for the Bokashi practitioners in the final workshop. (A) Microbial community composition in each stage, illustrated as Voronoi plots in which the polygon area correlates with the relative abundance of the genera. Top 10 genera are labelled in each plot. (B) Leachate characteristics. Organic acid concentrations are visualized in a bubble plot, in which the bubble size corresponds to the measured concentration in µM. Visualizations for all households are given in the supplementary material. Note that the original version of the visualization used during the workshop were prepared in Finnish language.

## Discussion

### Plant growth-promoting benefits of Bokashi compost application

Bokashi composting for the recycling of household food waste is increasing in popularity and comes with many promises, for many of which there is only anecdotal evidence reported by Bokashi practitioners. Open questions surrounding the practice of Bokashi, which also resonated strongly with the Bokashi practitioners participating in the present pilot study, include its useability as fertilizer, potential health risks through pathogens or potential health benefits through enrichment of beneficial microbes in the domestic ecosystem. The obtained results demonstrated the dominance of lactic acid bacteria in Bokashi ferment, and detected various short chain fatty acids such as lactate or acetate. In addition to the laboratory analyses, qualitative feedback from the study participants indicated a high perceived value of the ferment as a soil amendment. Even though these participant-reported outcomes were not quantitatively measured in this pilot, they provide experiential evidence of Bokashi’s functional potential in promoting plant growth. While this study is among the first that has studied Bokashi microbial communities on the household scale, more research has been done looking at the use of Bokashi for treatment of various waste materials on a larger scale as well as the use of the final product as an organic soil amendment in agriculture, and it is thus feasible to turn to those studies to look for answers to some of the study participants’ questions.

Microbial community composition in actively fermenting Bokashi has been only rarely assessed even on the agricultural scale, with slightly more studies looking at the microbial community composition in soils amended with Bokashi compost. Some studies have looked at the microbial community composition in Bokashi produced from animal manure, and the observed community composition is rather different from that observed in the present study: The community was more diverse and the most abundant taxa included Actinobacteriota, Proteobacteria and Chloroflexota (Abo-Sido *et al*. 2021; De Oliveira *et al*. 2025), and lactic acid bacteria were not a prominent part of the microbial communities. In comparison, relative abundances of Proteobacteria exceeded 10% only in half of the actively fermenting Bokashi samples in the present study, and Actinobacteria accounted for <2% of the microbial community in all actively fermenting Bokashi samples (Fig. 2). Bokashi compost has been used as an organic fertilizer e.g., in South America for a variety of crops including bananas, coffee, bell peppers, cucumber, tomatoes, maize and strawberries (Formowitz *et al*. 2007; Mosquera *et al*. 2016; Sarmiento, Amézquita and Mena 2019; Pandit *et al*. 2020; Abo-Sido *et al*. 2021; Mondaca *et al*. 2025; García-Hernández *et al*.). The addition of Bokashi compost increases crop production and soil fertility (e.g., (Gómez-Velasco *et al*. 2014; Jaramillo-López, Ramírez and Pérez-Salicrup 2015; Ferreira *et al*. 2017; Pandit *et al*. 2020)), and addition of biochar during or after the composting has an additional beneficial effect (Pandit *et al*. 2020; Pagliaccia *et al*. 2024; Dhakal *et al*. 2025). Observed benefits of Bokashi addition include i) increased nutrient availability and organic matter content (Gashua *et al*. 2023); ii) production of plant growth-promoting substances such as 3-phenyllactic acid (Maki *et al*. 2021), iii) increased microbial abundance (Gómez-Velasco *et al*. 2014; Mondaca *et al*. 2025), diversity (Luo *et al*. 2022), functional potential (Park and Kremer 2010) and activity (Gómez-Velasco *et al*. 2014) in soil, which might in turn accelerate nutrient turnover, iv) mitigation of the impact of heavy metal contamination (Barajas-Aceves and Rodríguez-Vázquez 2013), and v) increased suppressiveness of soils to plant pathogens such as fungi or root-knot nematodes in some cases (Ferreira *et al*. 2017; Shin *et al*. 2017; Van Der Sloot *et al*. 2024). These reported benefits appeal also to Bokashi practitioners on the household-scale, who often use their Bokashi compost as a soil amendment for gardening. However, waste material for Bokashi composting for agricultural use is more diverse than on the household-scale and often includes waste products such as chicken, pig or cow manure, fishery waste or even sludge from municipal wastewater treatment (e.g., (Epelde *et al*. 2018; Urra *et al*. 2019; Abo-Sido *et al*. 2021; De Oliveira *et al*. 2025)). These waste products typically have a higher nitrogen content than plant-based waste products, and the effect of plant-based Bokashi on plant growth might thus not be comparable, providing lower nitrogen amendment. On the other hand, application of organic amendments based on fermentation of animal manure of sewage can pose a higher risk of introduction of inorganic or organic contaminants, pathogens and AMR (Epelde *et al*. 2018; Urra *et al*. 2019), which is likely less problematic in household-scale fermentations using primarily plant-based waste products (with occasional animal products like fish bones or egg shells).

### Impact of Bokashi application on soil microbial communities and soil properties

The impact of soil amendment with Bokashi compost is inconclusive with some studies reporting changes in microbial community composition after addition of Bokashi fertilizer (Luo *et al*. 2022) and increases in bacterial CFU counts (Gashua *et al*. 2023), while others suggest no significant differences in bacterial community composition between Bokashi-amended and unamended soil, suggesting that Bokashi-microbes could not outcompete the native soil microbes (Shin *et al*. 2017). Indeed, microbial communities in soil factory samples in the present study strongly differed from starter, fermenting Bokashi and leachate samples (Fig. 2), and only the most abundant genus, *Lentilactobacillus*, was detected in three of the soil factories in notable abundances (Fig. 2B). The communities in the soil factories were rather similar to microbial communities reported from urban and suburban soils (e.g., (Baruch *et al*. 2021; Gómez-Brandón *et al*. 2022; Baldi *et al*. 2023; Probst *et al*. 2023; Tessler *et al*. 2023)). The most abundant genera detected in actively fermenting Bokashi, *Lentilactobacillus, Lactiplantibacillus, Companilactobacillus,* and *Levilactobacillus*, are homo- and heterofermentative lactic acid bacteria that are frequently found in fermented foods as well as food spoilage (Zheng *et al*. 2020). They can produce a variety of organic acids as well as ethanol. In the present study, lactate and acetate were the dominant organic acids produced, and fermentation pathways detected in metagenomic reads and MAGs confirmed the potential for diverse fermentations in actively fermenting Bokashi. Lactate, acetate and ethanol are also among the main fermentation products detected in Bokashi compost produced from plant-based waste on a larger scale (Pian *et al*. 2023). In addition, other studies have also reported the formation of butyrate (Alattar *et al*. 2012), which was likewise detected in the present study, albeit in low concentrations. Thus, while application of Bokashi compost to soils might not have a strong impact on the soil’s microbial community composition, it nonetheless adds organics acids, alcohols and other nutrients produced by lactic acid bacteria. Indeed, studies suggest that the benefits of Bokashi amendment are more likely due to the input of nutrients than the addition of living “effective” microorganisms to the soil (Schenck Zu Schweinsberg-Mickan and Müller 2009; Mayer *et al*. 2010; Kok *et al*. 2022). Bokashi addition does lead to a priming effect but not to long-term enhancement of soil organic carbon sequestration (Kok *et al*. 2022).

### Potential health benefits and risks of household-scale Bokashi composting

Use of Bokashi composting at the household scale features several routes of potential exposure to microorganisms from the Bokashi process: 1) inhalation or ingestion (e.g., after deposition on foods) of aerosolized microbes, 2) skin exposure during handling of Bokashi compost or soil factory, or 3) ingestion e.g., when using Bokashi as fertilizer for plants including fruits and vegetables. In general, lactic acid bacteria are known for health-promoting properties (Gilliland 1990), and many probiotics contain strains of lactic acid bacteria (Zielińska and Kolożyn-Krajewska 2018). Aerosolized lactic acid bacteria have been suggested as therapeutics to help combat lung diseases like infections with *P. aeruginosa* (Glieca *et al*. 2024), pulmonary neutrophilic inflammation (Nicola *et al*. 2024) or virus infections (Spacova *et al*. 2023). Bacteria and fungi can be aerosolized from household compost bins, but most of the aerosols are detected rather close to the compost bin (Naegele *et al*. 2016). Any negative effects (like occurrence of storage mites and molds) have been observed only close to the waste bin, not in the whole kitchen or home (Naegele *et al*. 2016), and it can be assumed that any potential positive impacts by aerosolized lactic acid bacteria would likewise be limited to the close proximity of the Bokashi bin or places in which leachate has been applied (e.g., flower pots). Study participants questioned whether direct skin contact with Bokashi during handling posed a risk of pathogen exposure. In larger settings, microbes from compost including lactic acid bacteria and potential pathogens have been detected on the hands of compost facility workers (Madsen, Rasmussen and Frederiksen 2024), which might indeed result in health risks if no proper hand hygiene is practiced. However, compost facilities receive waste from a large number of households, which greatly potentiates the risks compared to a single-household setting. In the present study, potential pathogens were only rarely detected in amplicons and metagenomes from Bokashi (0.005-0.5% of all reads), and most of them were opportunistic pathogens commonly found in soils or in the human microbiome. While risks for immuno-compromised people cannot be excluded, the pathogenic potential of Bokashi is fundamentally limited by its inputs. The presence of pathogens in the ferment likely reflects the existing household microbiota (e.g., from food waste, room air or skin contact), suggesting that the process does not introduce additional biological risks beyond those already present in the domestic environment. Among the non-bacterial species detected in the metagenomic reads in the present study was the parasitic whipworm *Trichuris trichiura*, which was detected in all samples (0.13-0.37% relative read abundance). Whipworm infections are not common in Europe (Behniafar *et al*. 2024), so it is highly unlikely that there has been contamination through a diseased person in the household. However, fruit and vegetable products imported from places where *T. trichiura* is endemic such as South America, Africa or East Asia could have had traces of said organism on the peel due to contamination (Alemu, Nega and Alemu 2020; Eslahi *et al*. 2022). *T. trichiura* has been detected on fruits and vegetables (e.g., banana, mango, cucumber, tomato) with a prevalence of 5-10% depending on the country of origin (Eslahi *et al*. 2022). Detection of DNA, however, does not give any indication whether there are actually living and/or infectious parasites present, as it might also persist in dead cells or as extracellular DNA.

Another concern is the introduction of AMR into the home environment. Indeed, lactic acid bacteria have been discussed as potential donors of AMR genes, and prospective probiotic food supplements have to undergo testing to prove that AMR transfer from these strains is not possible. Vancomycin resistance is intrinsic to many lactic acid bacteria like *Lactobacillus* and *Pediococcus* (Tynkkynen 1998), and genes encoding vancomycin resistance were detected in nearly all MAGs in the present study. However, as vancomycin resistance in lactic acid bacteria is chromosomally encoded and not closely related to enterococcal vancomycin resistance, it is believed not to be easily transferable (Tynkkynen 1998). Resistance to other antimicrobials like tetracycline, ciprofloxacin or chloramphenicol is fairly widespread in Lactobacilli with some carrying resistance to multiple antibiotics (Campedelli *et al*. 2019; Rivera-Rodriguez *et al*. 2025). Transfer of genes conveying resistance to streptomycin or erythromycin from Lactobacilli to other gram-positive and -negative species has been described (Gevers, Huys and Swings 2003; Ouoba, Lei and Jensen 2008; Tian *et al*. 2025). The MAGs recovered in the present study did encode a wide range of potential AMR genes and there might thus be a risk of AMR genes from Bokashi microbes to other species. However, it has to be kept in mind that after fermentation the Bokashi ferment was mixed with garden soil for the soil factory. Soils including garden soils are a huge reservoir of diverse AMR genes, and the interaction of humans with soils of any kind is thus associated with a certain risk (Popowska *et al*. 2012; Knapp *et al*. 2017; Lee *et al*. 2018; Zhao *et al*. 2025). It thus can be argued that handling the Bokashi ferment and using the soil from the soil factory does not carry a heightened risk compared to e.g., gardening.

### More questions than answers? – Recommendations for future studies

While this pilot study offers initial insights into microbial communities and experiential knowledge of urban Bokashi composting, it also highlights significant knowledge gaps. To address these gaps, future research should pursue two interlinked tracks: a participatory track (1) to ensure real-world relevance and a systematic track (2) to provide broader generalizability and statistical power.

(1) To align scientific inquiry with practitioner interests and needs, future projects should utilize co-design methodologies. Moving beyond passive participation, practitioners should be engaged as research partners from the outset. This collaborative framework would allow studies to address community-driven questions, such as the necessity of commercial inoculants or the perceived health benefits, ensuring that the research outputs are directly applicable by the Bokashi practitioners.
(2) To transition from pilot observations to generalized conclusions, upscaling and standardization are essential. Expanding the research to include larger, more diverse household cohorts will provide the statistical power required to account for the inherent variability of domestic waste. These efforts should be complemented by controlled laboratory experiments which allow for standardization of specific process variables (such as input food waste) that are difficult to control in a real household setting.

Based on the questions raised by this study’s participants, future research should prioritize three key areas: First, to address concerns regarding biosafety and potential health impacts, studies should quantify actual microbial exposure through bioaerosol monitoring and swabbing of skin household surfaces. Second, the benefits of Bokashi for plant growth require systematic testing to confirm (or refute) the practitioners’ subjective observations, and more detailed physicochemical analysis of Bokashi leachate and soil amendment is needed. Finally, starter optimization should be explored through comparative trials to determine whether commercial starter bran is strictly necessary or if it can be replaced by homemade microbial starters or inoculation with Bokashi ferment from previous batches. This would not only clarify the biological requirements of the process but also increase the affordability and long-term sustainability of household-scale Bokashi composting.

### Conclusions and limitations

Bokashi as a household-scale composting method suitable for urban lifestyle provides a promising low-technology, citizen-lead means to enhance microbial diversity and health of urban ecosystems. Bokashi practitioners, who have developed skills to make Bokashi with a sensory feeling for the process, achieve good fermentation results and productive soil without having a detailed microbiological understanding of the process. However, more information on the microbiological promises and risks of Bokashi are needed e.g., to allow health or environmental authorities to make recommendations or regulations concerning Bokashi composting. Bokashi composting on the household scale is surrounded by optimistic expectations to improve ecosystems and even human health, but the scientific basis for these claims is not well studied. In collaboration with Bokashi practitioners, this study assessed the microbial community composition in different stages of the Bokashi composting process. Lactic acid bacteria dominated starters and actively fermenting Bokashi, and metabolic pathways for the production of diverse organic acids were detected. While potentially pathogenic organisms and AMR genes were detected, their abundance was extremely low. Thus, handling Bokashi and keeping it in the house is likely not associated with increased health risks. Microbes from the Bokashi might be enriched in the household, e.g., due to aerosols near the compost bin. However, this was not assessed in the present study, and further studies are needed to assess the potential spread of microorganisms from the Bokashi compost in the household. Moreover, only a limited number of samples was analyzed in this study. This was in part due to limitations in the amount of funding, but also in part in the difficulty of recruiting suitable participants. Nonetheless, the study forms an important steppingstone for more detailed and more targeted studies on household-scale Bokashi composting.

## Supporting information

Supplemental Figutes

## Acknowledgements

The authors thank the Bokashi practitioners for active participation in the workshops and for providing samples as well as Sara Rinne for help with laboratory work.

## Author contributions

The article is based on interdisciplinary collaboration between Veera Kinnunen, who provided understanding of the lay Bokashi practice based on her ethnographic study, and Katharina Kujala, who contributed with microbiological expertise.

Katharina Kujala: Conceptualization (decision on sampling strategy, planning of physicochemical and molecular analyses), Formal analysis (bioinformatics), Investigation (laboratory work), Resources, Writing – Original draft, Writing – Review and editing, Visualization, Project administration, Funding acquisition

Veera Kinnunen: Conceptualization (General understanding of the lay Bokashi practice, planning and organization of participatory citizen workshops, participant recruitment), Investigation (leading discussions during workshops and summarizing input from participants), Resources, Writing – Review and editing, Project administration, Funding acquisition

## Funding

This work was supported by CitizenANTS funding provided by the Biodiverse Anthropocenes research program (supported by the University of Oulu and the Research Council of Finland PROFI6 funding (2021-2026), project Urbiolabs awarded to VK), the Research Council of Finland (project 322753 awarded to KK, and project number 362001 supporting VK) as well as the Jane and Aatos Erkko foundation (project ICE-FISHING awarded to KK).

## Data availability

The sequence data generated in this study has been deposited in the European Nucleotide Archive ENA under study accession number PRJEB100587.

## Notes

### Competing Interest Statement

The authors have declared no competing interest.

### Summary of Updates

Updated manuscript based on reviewer input from the journal it was submitted to.

