## Supplemental Figutes for "Lactic Acid Bacteria Dominate Urban Bokashi: A Participatory, Culture-independent Pilot Study of Microbial Diversity and Functional Potential in Household-Scale Food Waste Fermentation"

### Slide 1
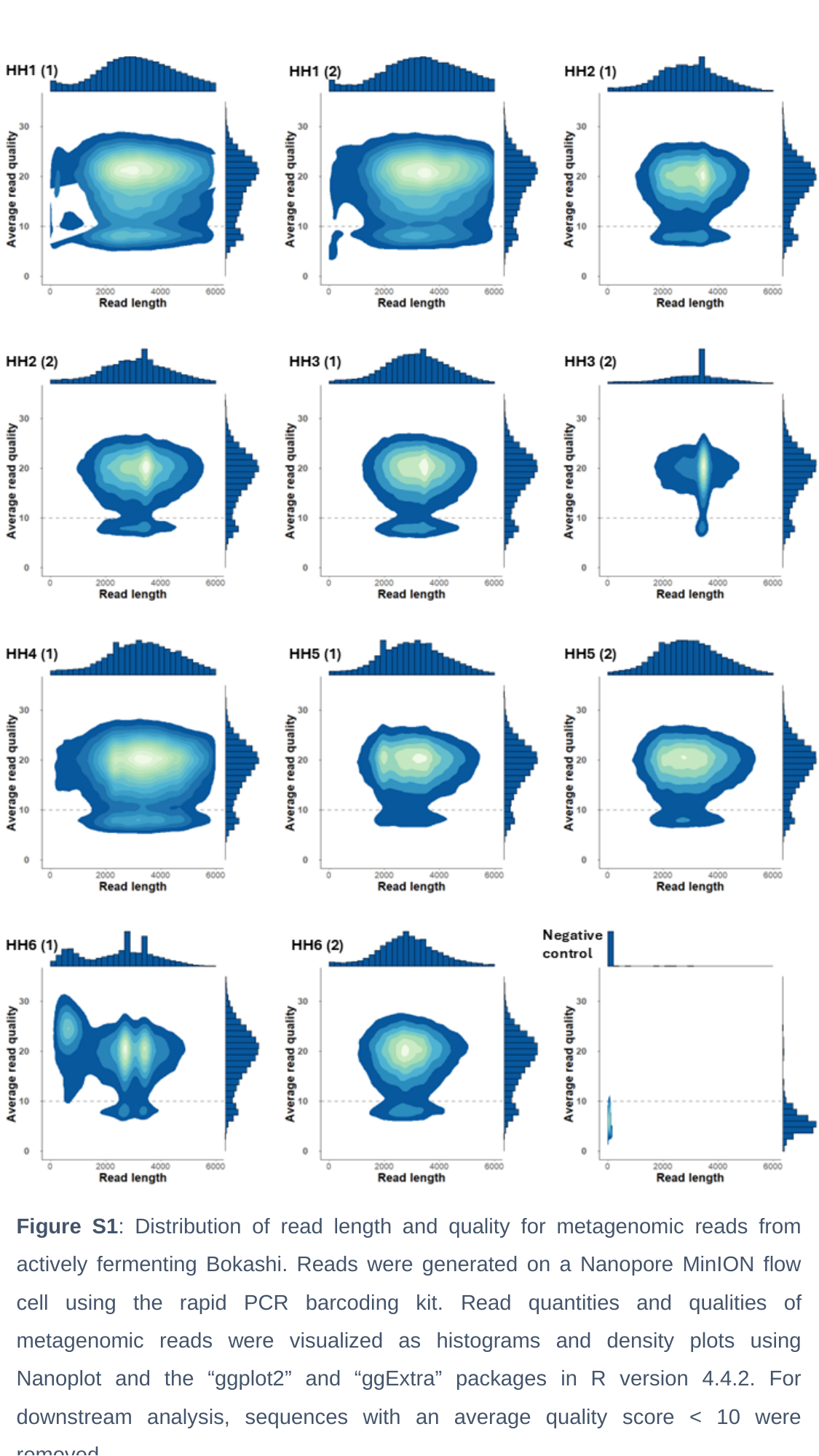

Figure S1: Distribution of read length and quality for metagenomic reads from actively fermenting Bokashi. Reads were generated on a Nanopore MinION flow cell using the rapid PCR barcoding kit. Read quantities and qualities of metagenomic reads were visualized as histograms and density plots using Nanoplot and the “ggplot2” and “ggExtra” packages in R version 4.4.2. For downstream analysis, sequences with an average quality score < 10 were removed.

### Slide 2
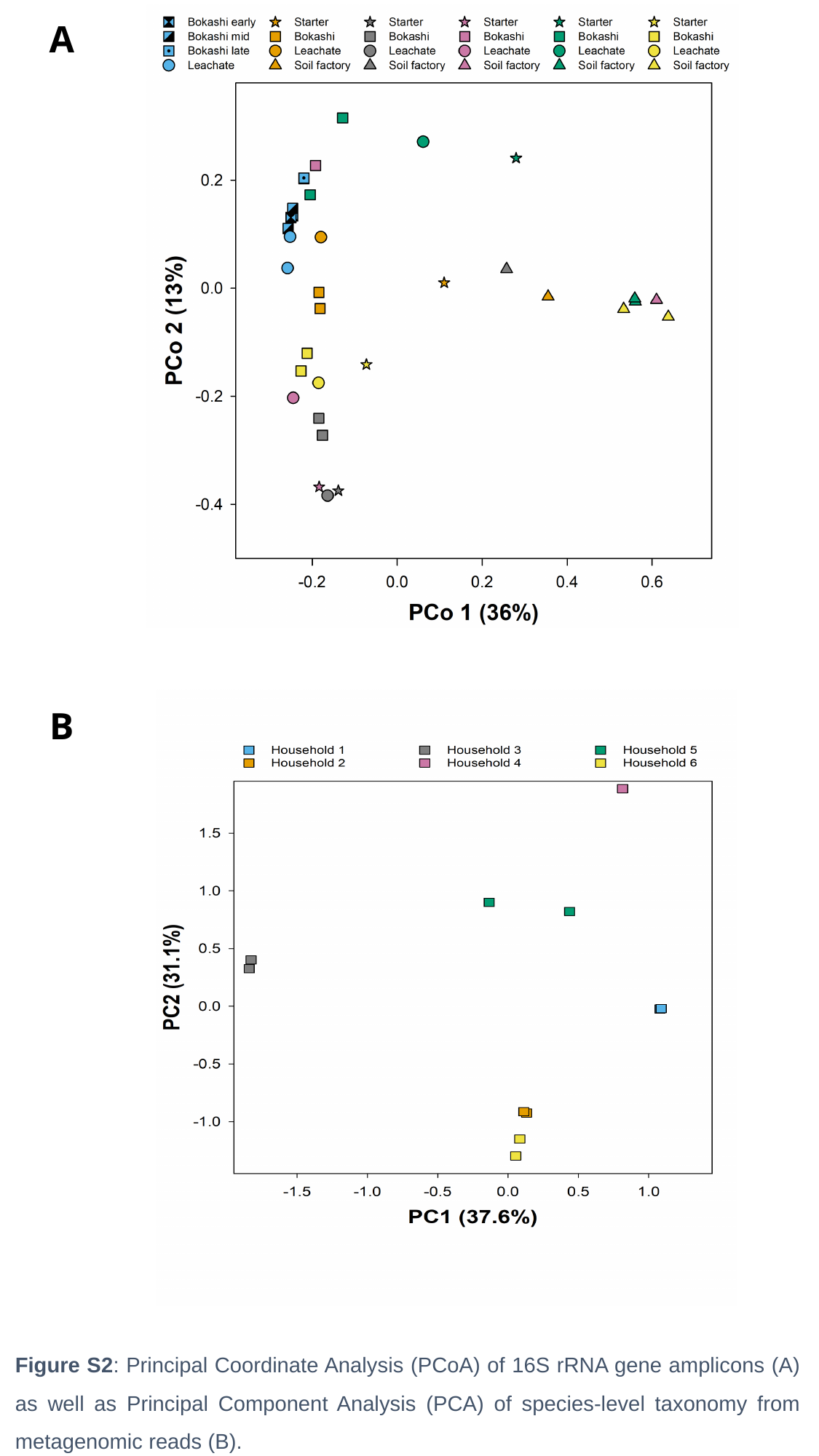

A
B
Figure S2: Principal Coordinate Analysis (PCoA) of 16S rRNA gene amplicons (A) as well as Principal Component Analysis (PCA) of species-level taxonomy from metagenomic reads (B).

### Slide 3
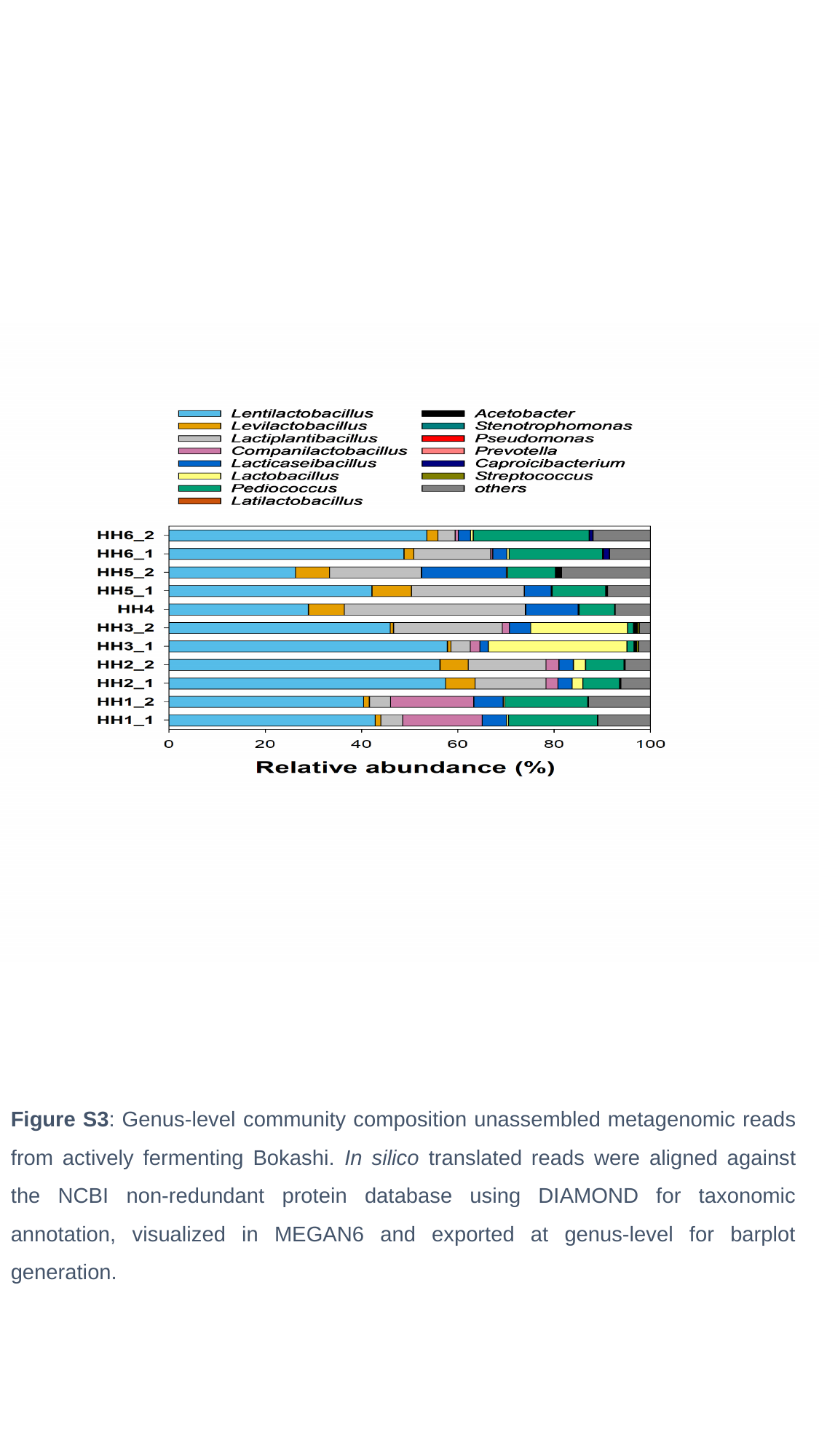

Figure S3: Genus-level community composition unassembled metagenomic reads from actively fermenting Bokashi. In silico translated reads were aligned against the NCBI non-redundant protein database using DIAMOND for taxonomic annotation, visualized in MEGAN6 and exported at genus-level for barplot generation.

### Slide 4
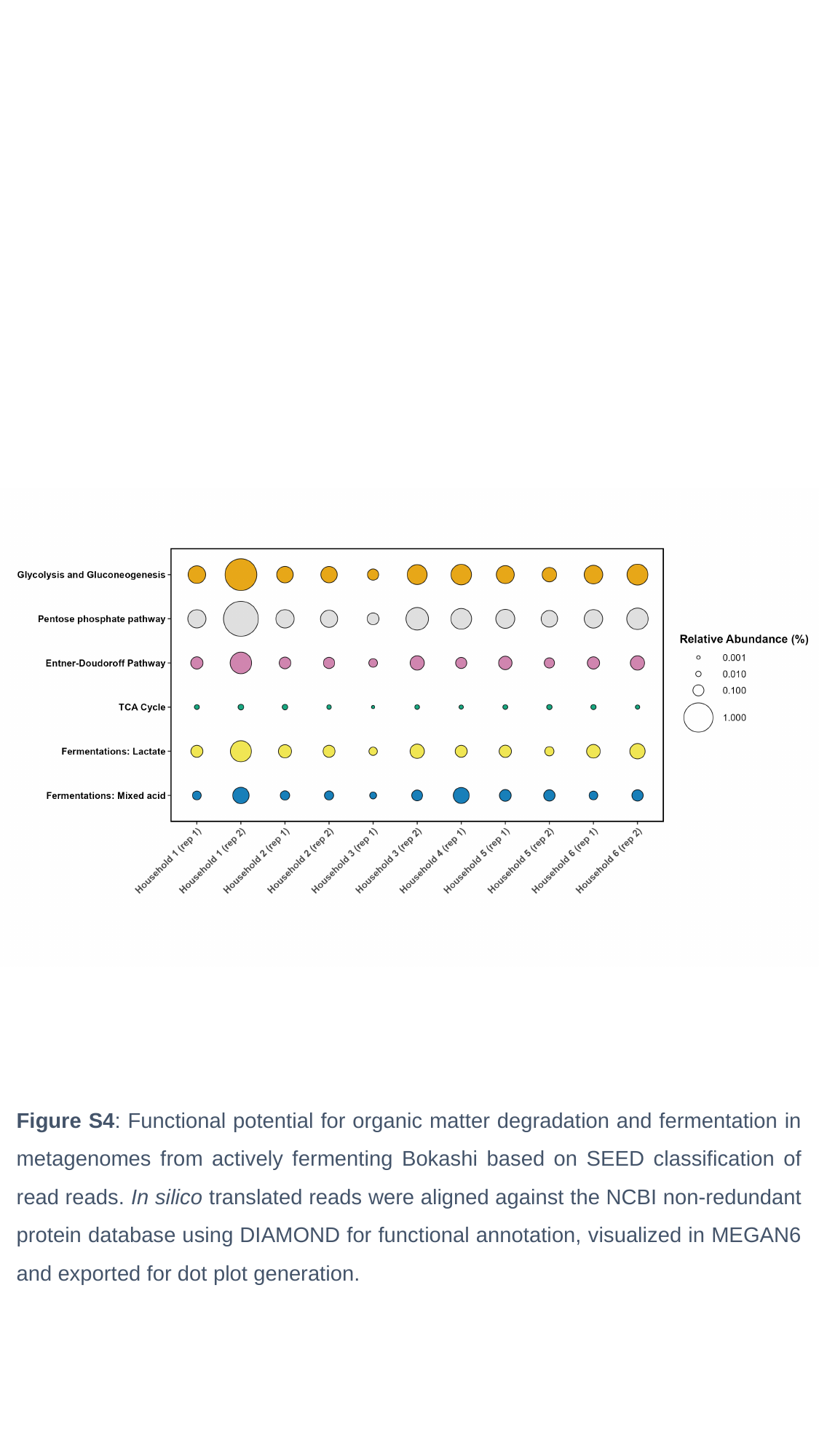

Figure S4: Functional potential for organic matter degradation and fermentation in metagenomes from actively fermenting Bokashi based on SEED classification of read reads. In silico translated reads were aligned against the NCBI non-redundant protein database using DIAMOND for functional annotation, visualized in MEGAN6 and exported for dot plot generation.

### Slide 5
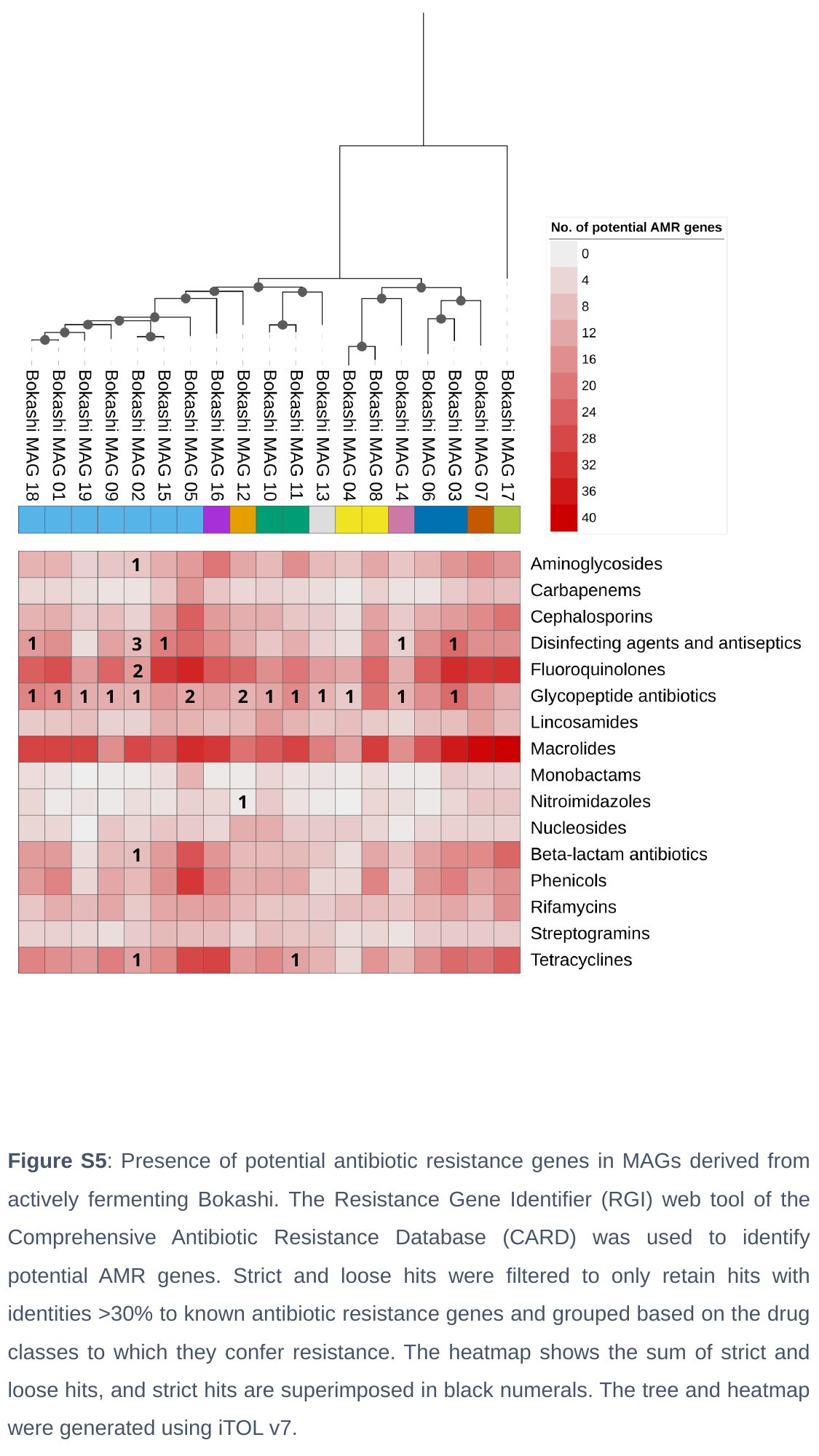

1
1
1
1
3
1
2
1
1
1
1
1
2
2
1
1
1
1
1
1
1
1
1
1
Figure S5: Presence of potential antibiotic resistance genes in MAGs derived from actively fermenting Bokashi. The Resistance Gene Identifier (RGI) web tool of the Comprehensive Antibiotic Resistance Database (CARD) was used to identify potential AMR genes. Strict and loose hits were filtered to only retain hits with identities >30% to known antibiotic resistance genes and grouped based on the drug classes to which they confer resistance. The heatmap shows the sum of strict and loose hits, and strict hits are superimposed in black numerals. The tree and heatmap were generated using iTOL v7.
